# Highly Multiplexed Immunofluorescence of the Human Kidney using Co-Detection by Indexing (CODEX)

**DOI:** 10.1101/2020.12.04.412429

**Authors:** Elizabeth K. Neumann, Emilio S. Rivera, Nathan Heath Patterson, Jamie L. Allen, Maya Brewer, Mark P. deCaestecker, Agnes B. Fogo, Richard M. Caprioli, Jeffrey M. Spraggins

**Affiliations:** Department of Biochemistry, Vanderbilt University, Nashville, TN, USA 37232; Mass Spectrometry Research Center, Vanderbilt University, Nashville, TN, USA 37232; Division of Nephrology and Hypertension, Department of Medicine, Vanderbilt University Medical Center, Nashville, TN USA 37232; Department of Pathology, Microbiology and Immunology, Vanderbilt University Medical Center, Nashville, TN USA 37232; Departments of Medicine and Pediatrics, Vanderbilt University Medical Center, Nashville, TN, USA 37232; Department of Chemistry, Vanderbilt University, Nashville, TN, USA 37232

**Keywords:** CODEX, Multiplexed Imaging, Immunofluorescence, Microscopy, Cell-type, Cell neighborhoods

## Abstract

The human kidney is composed of many cell types that vary in their abundance and distribution from organ to organ. As these cell types perform unique and essential functions, it is important to confidently label each within a single tissue to more accurately assess tissue architecture. Towards this goal, we demonstrate the use of co-detection by indexing (CODEX) multiplexed immunofluorescence for visualizing 23 antigens within the human kidney. Using CODEX, many of the major cell types and substructures, such as collecting ducts, glomeruli, and thick ascending limb, were visualized within a single tissue section. Of these antibodies, 19 were conjugated in-house, demonstrating the flexibility and utility of this approach for studying the human kidney using traditional antibody markers. We performed a pilot study showing that the studied tissues had on average 84 ± 11 cells per mm^2^ with the most variance seen within the cells containing vimentin and aquaporin 1, while cells containing α-smooth muscle actin and CD31 possessed a high degree of uniformity between the samples. These precursory data show the power of CODEX multiplexed IF for surveying the cellular diversity of the human kidney and have potential applications within pathology, histology, and building anatomical atlases.

---

The kidney is composed of over 20 cell types that can be further subdivided into discrete subpopulations, each performing an essential function for human health.^1,2^ Often, these cell types are implicated in the transition to disease states, such as diabetic nephropathy.^3–5^ Immunohistochemical approaches are the standard for labeling cell types and structures within a tissue matrix because of their inherent specificity, application to a wide variety of protein targets, and insight into biochemical pathways.^6,7^ Fluorescently tagged antibodies are often used because they provide low background for high signal-to-noise imaging at spatial resolutions between 250 nm to 1 μm.^8,9^ Fluorescence experiments, however, are limited in plexity to ~4-7 antibodies because of spectral overlap between commonly used fluorophores.^10–12^ Traditional cyclic multiplexed immunofluorescence approaches consist of antibody application, imaging, and fluorescence inactivation/bleaching before starting a new cycle. These methods increase the number of imageable targets between 20-60 but are ultimately limited by tissue distortion and antibody-antibody interactions.^13^

Recently, co-detection by indexing (CODEX) multiplexed immunofluorescence (IF) was developed to improve multiplexed imaging efforts to over 50 antigens.^14,15^ Briefly, oligonucleotide barcodes are conjugated to primary antibodies and all primary antibodies are incubated and fixed to either fresh frozen or paraffin embedded tissue. Complementary oligonucleotide barcodes attached to fluorophores are serially added, imaged, and removed to build highly multiplexed imaging datasets without significant tissue degradation. Here, we demonstrate the power of CODEX IF to image 23 cell-defining antigens within multiple human kidneys using a mixture of commercial and in-house conjugated antibodies. This multiplexed IF workflow has comparable or higher plexity than other multiplexed imaging approaches produced to date and can image larger tissue areas (on the order of mm^2^) than multiplexed technologies such as cyTOF,^16,17^ demonstrating the potential of this technology for exploring cellular heterogeneity within the human kidney.

## RESULTS

### Multiplexed IF Imaging using CODEX

We have generated CODEX multiplexed IF images from human kidney tissue using 23 barcoded antibodies (Table 1, SI Table 1). The major structures within the kidney were imaged, such as glomeruli, tubules, and collecting ducts by targeting CD93, aquaporin 1, and cytokeratin 7, respectively. These structures are further visually subdivided by targeting antigens that localize to different tubular layers (e.g. CD90, β-catenin, E-cadherin, and calbindin). Many of these selected antigens assist in defining important cell types and have been previously studied by standard immunofluorescence studies, allowing comparison of our novel approach with previous results. When possible, we selected commercial, recombinant, and purified primary antibodies for conjugation. These criteria enable reproducible staining, limiting the commonly cited weaknesses of antibody-based approaches (e.g. irreproducible staining and lot-to-lot variability).^18,19^ The tissue remained intact through the eleven cycles performed in this study (two blank cycles and nine antibody-containing cycles), indicating that additional cycles could even be incorporated to increase the number of antibody targets further.

**Table 1:** Summary of the cell types visualized by CODEX multiplexed IF. Italicized rows indicate commercial CODEX markers. Other antibodies were conjugated to CODEX oligonucleotide barcodes by us.

| Antigen | Cell Type | Barcode | Channel | Cycle |
| --- | --- | --- | --- | --- |
| *CD7* | *Natural killer cells, T-cells* | *BX025* | *FITC* | *1* |
| Aquaporin 1* | Proximal tubule cells, endothelial cells, thin descending limb of the loop of Henle | BX002 | DsRed | 1 |
| Aquaporin 2* | Principal cells of the collecting duct | BX015 | Cy5 | 1 |
| *CD90* | *Proximal tubules, fibroblasts, activated endothelial cells* | *BX022* | *FITC* | *2* |
| CD38* | T cells, B cells, NK cells, erythrocytes | BX007 | FITC | 3 |
| *Ki67* | *Proliferating cells* | *BX026* | *DsRed* | *3* |
| Nestin* | Podocytes, endothelial cells | BX024 | Cy5 | 3 |
| *CD45* | *All hematopoietic cells, except erythrocytes and platelets* | *BX001* | *FITC* | *4* |
| Uromodulin* | Epithelial cells of the thick ascending limb of Henle's loop | BX047 | DsRed | 4 |
| β-Catenin* | Tubular epithelium | BX003 | Cy5 | 4 |
| Vimentin* | Mesenchymal cells, tubular cells, glomerular epithelial cells | BX043 | Cy5 | 5 |
| PARP1* | Glomerular endothelium | BX013 | FITC | 6 |
| *CD31* | *Endothelial cells* | *BX032* | *DsRed* | *6* |
| CD93* | Peritubular capillary and glomerular endothelium | BX006 | Cy5 | 6 |
| E-Cadherin | Distal convoluted tubules, collecting duct, loop of Henle | BX016 | FITC | 7 |
| Laminin, gamma-1 | Basement membrane | BX023 | DsRed | 7 |
| KDR/Vegf * | Peritubular capillary and glomerular endothelium | BX030 | Cy5 | 7 |
| Calbindin* | Distal convoluted tubules, connecting segment, and cortical collecting duct | BX004 | FITC | 8 |
| Cytokeratin 7* | Epithelial cells | BX005 | DsRed | 8 |
| α-Smooth Muscle Actin* | Vascular smooth muscle, myofibroblasts | BX021 | Cy5 | 8 |
| Renin* | Juxtaglomerular cell | BX010 | FITC | 9 |
| Tryptase* | Mast cells | BX014 | DsRed | 9 |
| MARCKS* | Peritubular capillary | BX027 | Cy5 | 9 |

CODEX imaging was performed on three normal non-neoplastic portions of nephrectomy samples, two men and one woman, 47, 66, and 77 years old, respectively (Figure 1, SI Figures 1 and 2). Images from large sections were acquired (Figure 1A), including areas of cortex and medulla, with application of several antibodies binding exclusively to cells in the medulla (Figure 1B) or cortex (Figure 1C). The distribution of interstitial cells was also visualized, such as mast cells. Special markers of the juxtaglomerular region of the nephron, key for nephron hemodynamic responses, were also assessed. Further, different cell types and regions within a single glomerulus were also resolved (SI Figure 3), specifically, the glomerular endothelium, podocytes, and basement membrane. The data demonstrate the unique distribution of these cells within the glomeruli as visualized using PARP1, calbindin, laminin, and nestin.

**Figure 1:**
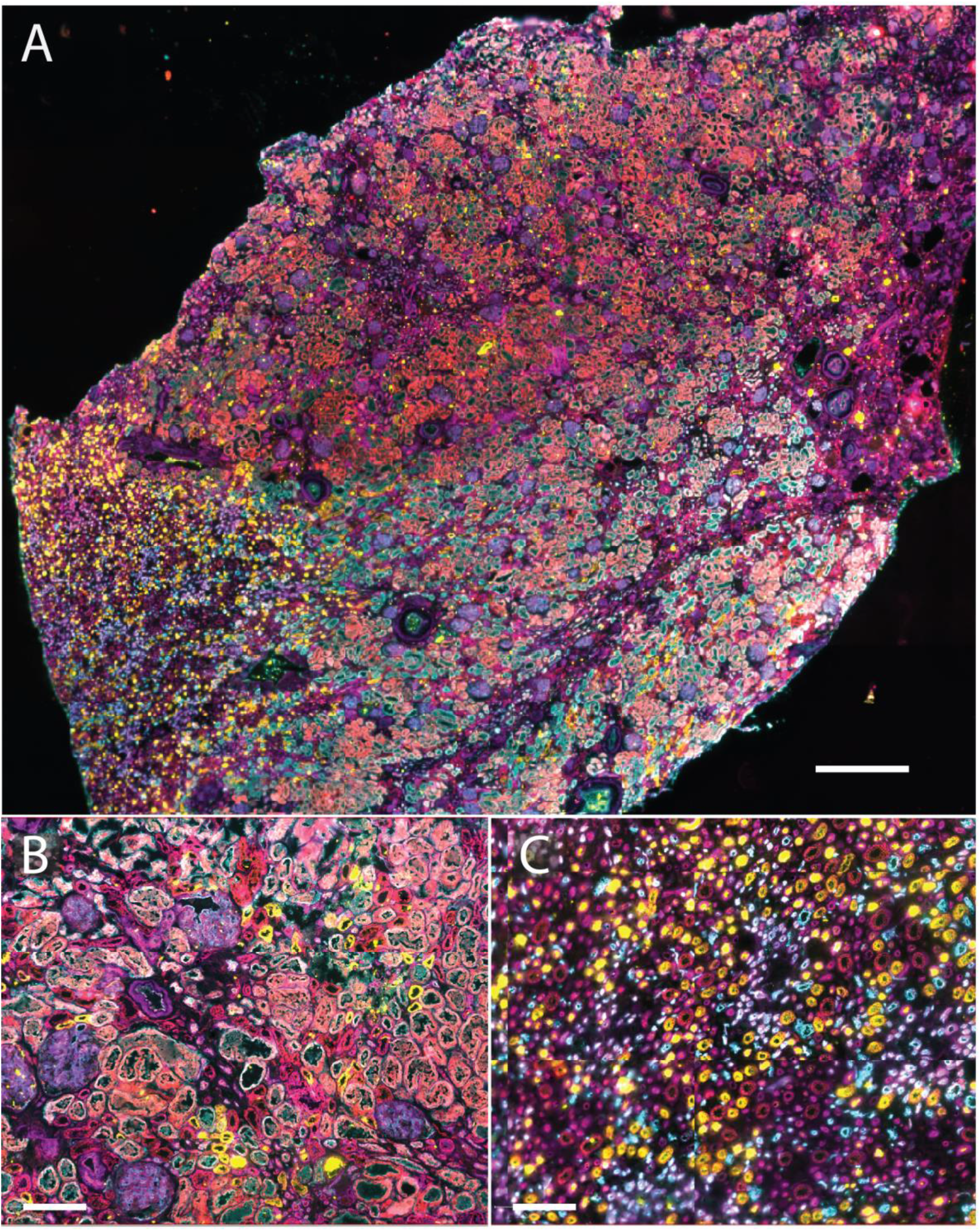
**A)** CODEX multiplexed IF staining of the human kidney. Scale bar 1 mm. **B)** Enlargement of a region of the cortex with several glomeruli present as well as proximal tubules. Scale bar 200 μm. **C)** Medullary region of the kidney with many tubules present. Scale bar 200 μm. The color legend for all panels is as follows: cytokeratin 7 (pink), α-smooth muscle actin (red), renin (dark pink), uromodulin (yellow), ß-catenin (red), aquaporin 1 (teal), KDR (dark red).

### Cell Type and Neighborhood Analysis

To show the potential of highly multiplexed molecular analysis, we calculated the number of each cell type per tissue area (Figure 2). On average, the tissue contained 84 ± 11 cells/mm^2^. This analysis can be extended to include specific cell subtypes. Cells with high levels of α-smooth muscle actin were consistently present in all samples, while in contrast, vimentin-containing cells were the most variable cell type (Figure 2A). Unsurprising, when adjusted for medulla and cortex content, variability between the patients decreased (Figure 2B). Finally, we also could assess how cells are spatially organized within the tissue (Figure 2C). For instance, cells expressing CD38 are often neighbors with cells expressing ki67 (correlation score>2), whereas cells expressing laminin are distant from cells expressing α-smooth muscle actin (correlation score <-2). While only some examples are discussed here, similar comparisons can be made for all the antigens stained using CODEX. These preliminary analyses importantly demonstrate the ability to survey cell populations with high sensitivity and selectivity using multiplexed molecular imaging and the potential of this technique to illuminate renal structure and function.

**Figure 2:**
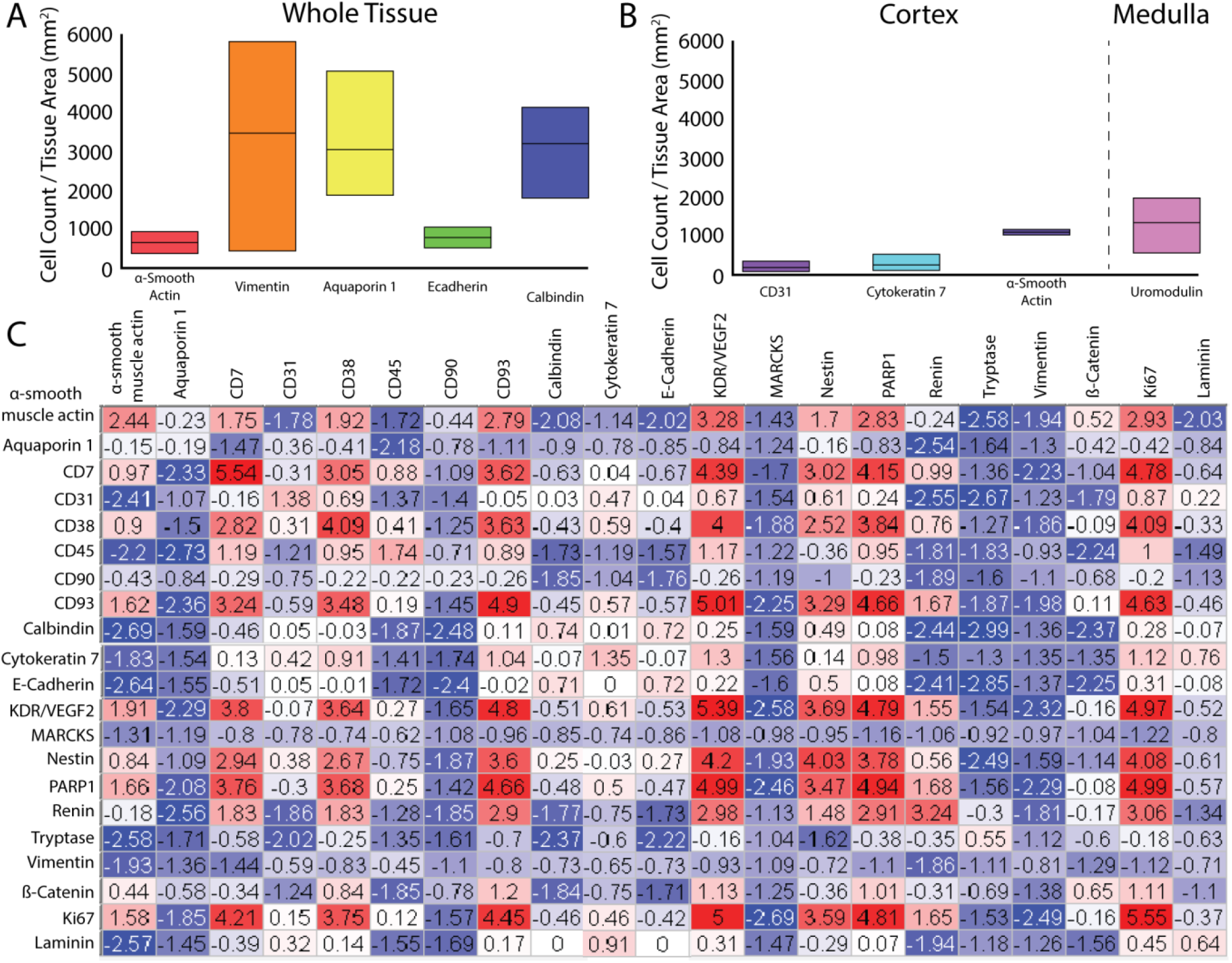
**A)** Average cell count normalized by tissue area. Highest variability is seen in the amount of vimentin and lowest in e-cadherin. **B)** Average cell count normalized by tissue subregion based on highest abundance. CD31, cytokeratin 7, and α-smooth muscle actin were detected primarily in the cortex and uromodulin was detected primarily in the medulla. **C)** Correlation plot from one kidney sample indicating cell types that are highly correlated (red), noncorrelated (white), and uncorrelated (blue).

## DISCUSSION

Using 23 antibodies, key renal cell types can be visualized and compared in human kidney tissue. This study incorporates a combination of commercial CODEX labels as well as conjugated purified antibodies from three different vendors, demonstrating the flexibility of the approach to accommodate a variety of antibody sources. Our studies support that well-validated, primary antibodies free from common preservations (e.g. bovine serum albumin, glycerol, and sodium azide) are preferable for this approach. Once proper primary antibodies are selected and conjugated, it is important to validate conjugation does not changing the binding efficiency or selectivity of the original primary antibody (SI Figure 4). Additionally, selecting recombinant antibodies reduces variability and is essential for creating large renal atlases, such as those pioneered by the Kidney Precision Medicine Project and Human BioMolecular Atlas Program, which require examination of renal sections from many patients over the course of years. Most of the antibodies used here are recombinant (Table 1, *) and can serve as an initial panel for applying CODEX IF to fresh frozen human kidney tissues or biopsies for these critical atlas efforts. Moreover, we anticipate that study of kidney diseases will also benefit from the development of similar multiplexed panels that may aid in assessment of complex cell interplay and pathophysiology of a spectrum of injury. Diseases with similar light microscopic appearances may have varying phenotypes and alterations of specific cells, which may shed light on mechanisms and potential targets for intervention. While this study was performed on fresh frozen tissue, many of these antibodies are validated for formalin-fixed paraffin embedded (FFPE) sections, thus allowing for application of CODEX IF to FFPE sections. Such highly multiplexed analysis could pave the way for detailed phenotyping, classification and understanding of diseases, enhancing research and patient care.

## METHODS

Normal portions of renal cancer nephrectomies from adult patients were studied. Tissue blocks were collected from areas distant to the tumor, frozen over an isopentane dry ice slurry, and 10 μm sections were thaw-mounted onto glass cover slips. These sections were then fixed and incubated with a mixture containing all primary antibodies. Secondary oligonucleotide sequences were automatically added and removed serially for multiplexed visualization on a single section without spectral overlap using the commercial Akoya platform (Akoya Biosciences, Marlborough, MA). Resulting images were background subtracted, cycle aligned, processed using extended depth of field, and stitched using the CODEX processor and MAV software (Akoya Biosciences). Water shed cell segmentation was performed using MAV software. Fluorescence intensity greater than background was used as a positive label. Full images were interactively visualized in QuPath after conversion to the pyramidal OME.TIFF open standard format.^20^ Further experimental details are in the Supplemental Information.

## ACKNOWLEGEMENTS

Support was provided by the NIH Common Fund and National Institute of Diabetes and Digestive and Kidney Diseases (NIDDK) (U54DK120058 awarded to J.M.S. and R.M.C.) and NIH National Institute of Allergy and Infectious Disease (NIAID) (R01AI138581 awarded to J.M.S.). E.K.N. is supported by a National Institute of Environmental Health Sciences training grant (T32ES007028). Human tissues were acquired through the Cooperative Human Tissue Network at Vanderbilt University Medical Center which is supported by the NIH National Cancer Institute (5 UM1 CA183727-08).

## REFERENCES

(1) Al-Awqati, Q.; Oliver, J. A. Kidney International 2002, 61, 387.

(2) Puelles, V. G.; Hoy, W. E.; Hughson, M. D.; Diouf, B.; Douglas-Denton, R. N.; Bertram, J. F. Current Opinion in Nephrology and Hypertension 2011, 20.

(3) Gambara, V.; Mecca, G.; Remuzzi, G.; Bertani, T. Journal of the American Society of Nephrology 1993, 3, 1458.

(4) Toyota, E.; Ogasawara, Y.; Fujimoto, K.; Kajita, T.; Shigeto, F.; Asano, T.; Watanabe, N.; Kajiya, F. Kidney International 2004, 66, 855.

(5) Najafian, B.; Alpers, C. E.; Fogo, A. B. Contrib Nephrol 2011, 170, 36.

(6) Buchwalow, I. B.; Böcker, W. Basics and Methods 2010, 1, 1.

(7) Duraiyan, J.; Govindarajan, R.; Kaliyappan, K.; Palanisamy, M. Journal of pharmacy & bioallied sciences 2012, 4, S307.

(8) Hell, S. W.; Dyba, M.; Jakobs, S. Current Opinion in Neurobiology 2004, 14, 599.

(9) Bastiaens, P. I. H.; Squire, A. Trends in Cell Biology 1999, 9, 48.

(10) Schubert, W.; Bonnekoh, B.; Pommer, A. J.; Philipsen, L.; Böckelmann, R.; Malykh, Y.; Gollnick, H.; Friedenberger, M.; Bode, M.; Dress, A. W. M. Nature Biotechnology 2006, 24, 1270.

(11) Gerdes, M. J.; Sevinsky, C. J.; Sood, A.; Adak, S.; Bello, M. O.; Bordwell, A.; Can, A.; Corwin, A.; Dinn, S.; Filkins, R. J.; Hollman, D.; Kamath, V.; Kaanumalle, S.; Kenny, K.; Larsen, M.; Lazare, M.; Li, Q.; Lowes, C.; McCulloch, C. C.; McDonough, E.; Montalto, M. C.; Pang, Z.; Rittscher, J.; Santamaria-Pang, A.; Sarachan, B. D.; Seel, M. L.; Seppo, A.; Shaikh, K.; Sui, Y.; Zhang, J.; Ginty, F. Proceedings of the National Academy of Sciences 2013, 110, 11982.

(12) Tsurui, H.; Nishimura, H.; Hattori, S.; Hirose, S.; Okumura, K.; Shirai, T. Journal of Histochemistry & Cytochemistry 2000, 48, 653.

(13) Lin, J.-R.; Izar, B.; Wang, S.; Yapp, C.; Mei, S.; Shah, P. M.; Santagata, S.; Sorger, P. K. eLife 2018, 7, e31657.

(14) Goltsev, Y.; Samusik, N.; Kennedy-Darling, J.; Bhate, S.; Hale, M.; Vazquez, G.; Black, S.; Nolan, G. P. Cell 2018, 174, 968.

(15) Park, J.; Liu, C.; Kim, J.; Susztak, K. Kidney International 2019, 96, 862.

(16) Fribourg, M.; Anderson, L.; Fischman, C.; Cantarelli, C.; Perin, L.; La Manna, G.; Rahman, A.; Burrell, B. E.; Heeger, P. S.; Cravedi, P. Kidney International 2019, 96, 436.

(17) Singh, N.; Avigan, Z. M.; Kliegel, J. A.; Shuch, B. M.; Montgomery, R. R.; Moeckel, G. W.; Cantley, L. G. JCI Insight 2019, 4, e129477.

(18) Jarvis, M. F.; Williams, M. Trends in Pharmacological Sciences 2016, 37, 290.

(19) Gong, B.; Murray, K. D.; Trimmer, J. S. New Biotechnology 2016, 33, 551.

(20) Bankhead, P.; Loughrey, M. B.; Fernández, J. A.; Dombrowski, Y.; McArt, D. G.; Dunne, P. D.; McQuaid, S.; Gray, R. T.; Murray, L. J.; Coleman, H. G.; James, J. A.; Salto-Tellez, M.; Hamilton, P. W. Scientific Reports 2017, 7, 16878.

