## Supplemental Information for "Highly Multiplexed Immunofluorescence of the Human Kidney using Co-Detection by Indexing (CODEX)"

### SUPPORTING INFORMATION

#### Table of Contents

Table S1 Antibody Table ........ S-2

Figure S1 CODEX IF Image of Second Patient ........ S-3

Figure S2 CODEX IF Image of Third Patient ........ S-4

Figure S3 CODEX IF Images of Glomeruli ........ S-4

Figure S4 Comparison between Indirect IF and CODEX Conjugated Antibodies ........ S-5

Extended Methods ........ S-5

| Antigen | Primary Antibody Product Code | Barcode |
|---|---|---|
| CD7 | Akoya Biosciences 4150022 | *BX025* |
| Aquaporin 1 | Abcam b9566 | BX002 |
| Aquaporin 2 | Abcam ab230170 | BX015 |
| CD90 | Akoya Biosciences 4150021 | *BX022* |
| CD38 | Akoya Biosciences 4150007 | BX007 |
| Ki67 | Akoya Biosciences 4250019 | *BX026* |
| Nestin | Abcam ab221660 | BX024 |
| CD45 | Akoya Biosciences 4150003 | *BX001* |
| Uromodulin | Abcam ab207170 | BX047 |
| ß-Catenin | Abcam ab32572 | BX003 |
| Vimentin | Abcam ab92548 | BX043 |
| PARP1 | Abcam ab191217 | BX013 |
| CD31 | Akoya Biosciences 4250009 | *BX032* |
| CD93 | Abcam ab134079 | BX006 |
| E-Cadherin | Cell Signaling 3195S | BX016 |
| Laminin, gamma-1 | DSHB 2E8 | BX023 |
| KDR/Vegf 2 | Abcam ab237634 | BX030 |
| Calbindin | Abcam ab108404 | BX004 |
| Cytokeratin 7 | Abcam ab68459 | BX005 |
| α-Smooth Muscle Actin | Abcam ab7817 | BX021 |
| Renin | Abcam ab212197 | BX010 |
| Tryptase | Abcam ab151757 | BX014 |
| MARCKS | Abcam ab184546 | BX027 |

**Table S1:** Summary of the cell types visualized by CODEX multiplexed IF with product information.

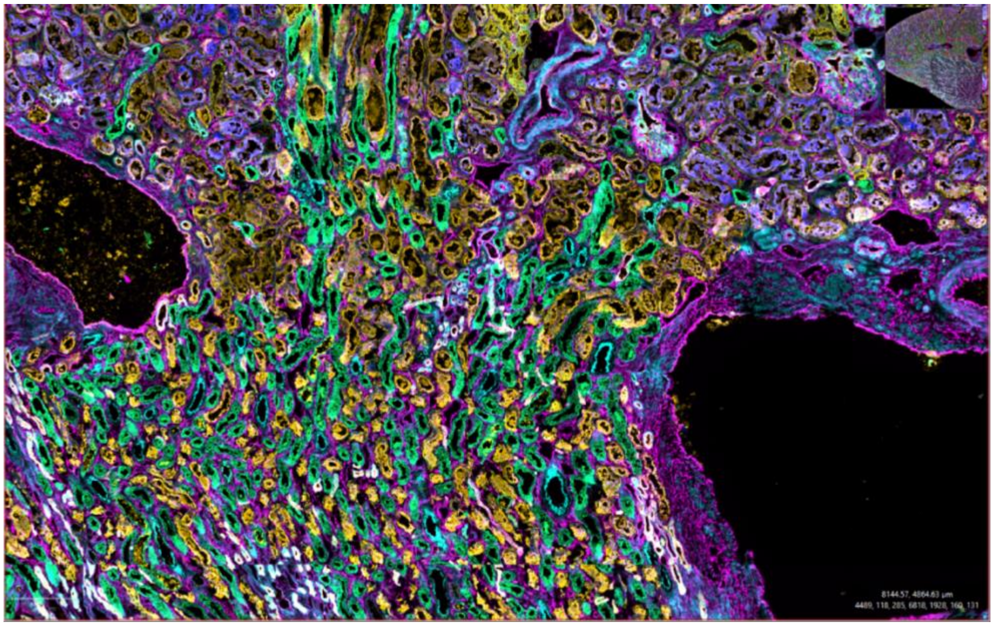


**Figure S1:** CODEX multiplexed immunofluorescence of kidney section for a 77-year-old female, using cytokeratin 7 (yellow), α-smooth muscle actin (blue), vimentin (pink), aquaporin 1 (dark yellow), aquaporin 2(teak), uromodulin (light green).

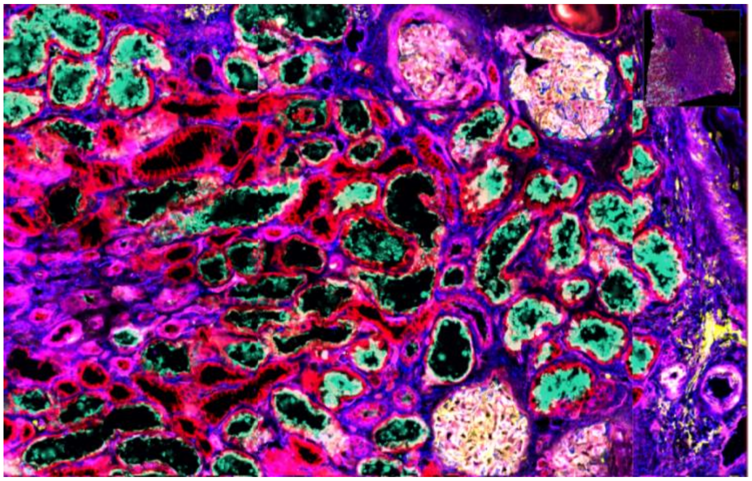

**Figure S2:** CODEX multiplexed immunofluorescence of kidney section for a 44-year-old female, using ß-catenin (red), vimentin (blue), VEGF 2 (red), calbindin (yellow), cytokeratin 7 (pink), α-smooth muscle actin (teal).

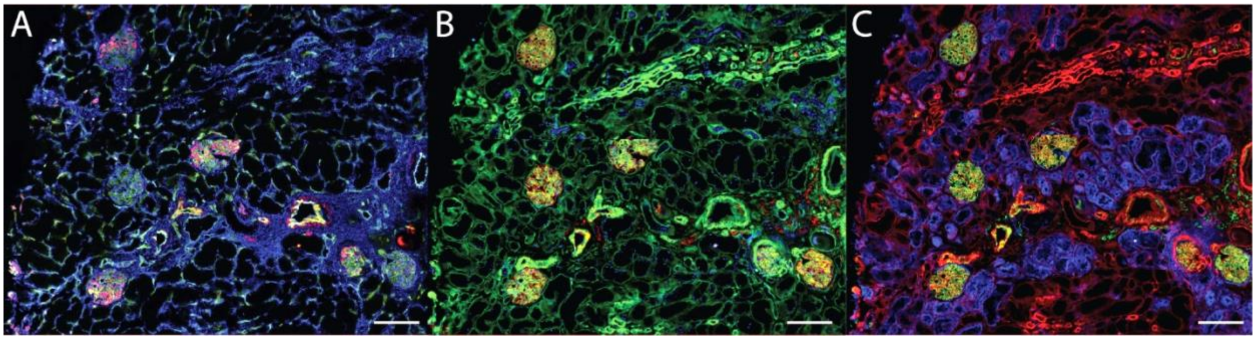


**Figure S3:** CODEX multiplexed immunofluorescence visualization of glomeruli, using **A)** nestin (red), vimentin (blue), CD31 (green) **B)** PARP1 (blue), e-cadherin (red), laminin (green), and **C)** calbindin (green), cytokeratin 7 (red, CD90 (blue).

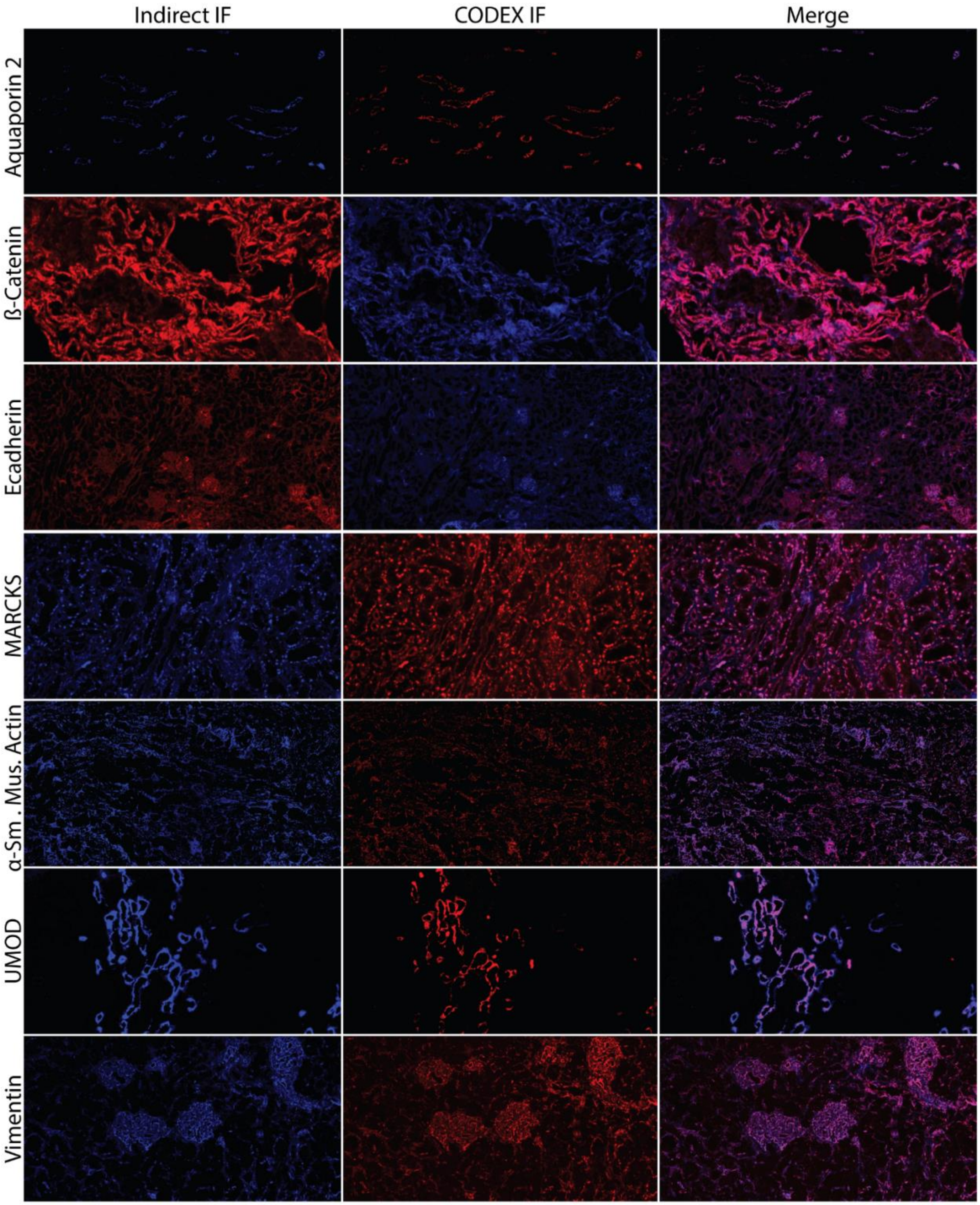


**Figure S4:** Comparison of indirect IF versus CODEX conjugated antibodies using the same primary antibody. Overlaid images show high concordance between the indirect IF and CODEX IF.

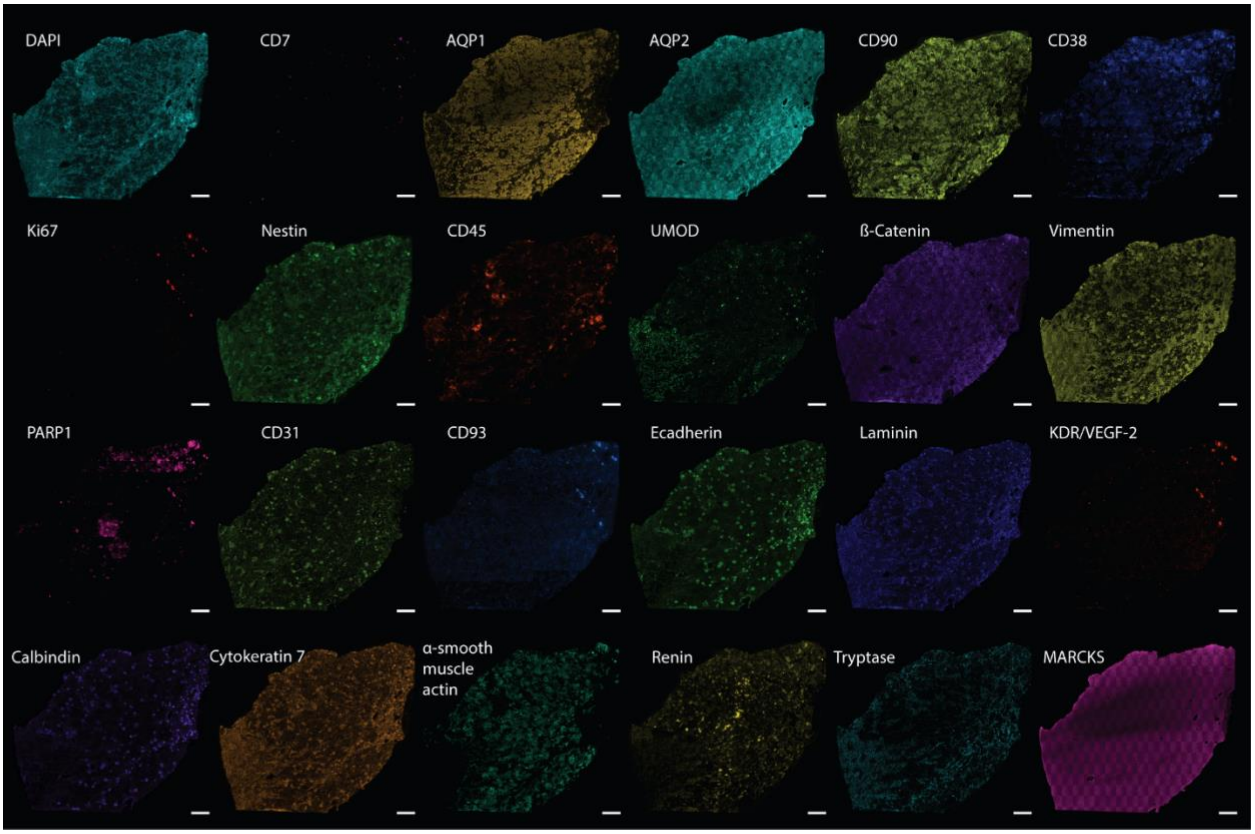


**Figure S5:** Single channel images of CODEX IF markers.

**Extended Methods:**

**Materials:**

HPLC-grade acetone, isopentane, and methanol were purchased from Fisher Scientific (Pittsburgh, PA, USA).

**Sample Preparation:**

Human kidney tissue was surgically removed during a full nephrectomy and remnant tissue was processed for research purposes by the Cooperative Human Tissue Network at Vanderbilt University Medical Center. Remnant biospecimens were collected in compliance with the Cooperative Human Tissue Network standard protocols and National Cancer Institute's Best Practices for the procurement of remnant surgical research material. Participants were consented for remnant tissue collection in accordance to institutional IRB policies. The excised tissue was flash frozen over an isopentane-dry ice slurry, embedded in carboxymethylcellulose, and stored at -80 °C until use. Kidney tissues were cryosectioned to a 10 µm thickness and thaw mounted onto poly-L-lysine coated glass cover slips and stored at -80 °C until analyzed.

Tissues were returned to ~20 °C within a vacuum desiccator before being prepared according to the manufacturer's protocols (Akoya Biosciences, Marlborough, MA). In brief, tissues were rehydrated in hydration buffer (Akoya Biosciences) and fixed for 10 minutes in a 1.6% paraformaldehyde (Thermo Fischer Scientific, Waltham, MA) solution diluted in hydration buffer. Tissues were incubated in staining buffer (Akoya Biosciences) for 30 min. Tissues were then incubated with blocking buffer (Akoya Biosciences) and primary antibody cocktail (1:200 antibody dilution) for 3 hr within a humidity chamber at ~20 °C. The samples were rinsed with staining buffer before being fixed in a 1.6% paraformaldehyde solution diluted in storage buffer (Akoya Biosciences) for 10 min. The samples were then rinsed in phosphate buffered saline (PBS), incubated for 5 min in cold methanol (~4 °C), and rinsed again in PBS. Samples were then fixed with fixative solution (Akoya Biosciences) before being stored in storage buffer at ~4 °C until used.

**Antibody Conjugation:**

Pre-conjugated CODEX antibodies were purchased from Akoya Biosciences. Unconjugated primary antibodies were purchased from Abcam (Cambridge, MA) unless otherwise specified (SI Table 1). Purified primary antibodies were purchased without any additives or preservatives, many of which prevent successful antibody conjugation. Antibodies were conjugated and prepared according to the manufacturer's protocols (Akoya Biosciences) with slight deviations. 50 kDa molecular weight cut off filters were blocked using 500 µL of a proprietary filter blocking solution (Akoya Biosciences). 50 µg of each antibody were diluted to 100 µL of PBS and filtered. Antibodies were reduced with the reduction mixture (Akoya Biosciences) for 25 min at ~20 °C. The reduction solution was removed by centrifugation (12,000 g) and exchanged with buffer solution (Akoya Biosciences). Each oligonucleotide barcode is rehydrated in 10 µL nuclease free water (Ambion Inc., Austin, TX) and further diluted in 210 µL of conjugation solution (Akoya Biosciences). Respective barcodes are added to the reduced primary antibodies and incubated for 2 hr at ~20 °C. 5 µL of this conjugated antibody solution is removed for later validation. The solution is than purified by buffer exchanging with purification solution (Akoya Biosciences) and stored in storage solution (Akoya Biosciences) at ~4 °C until used. After verifying conjugation, the staining profiles of newly conjugated antibodies were compared to a traditional, indirect immunofluorescence experiment.

**CODEX Multiplexed Immunofluorescence:**

Antibodies are diluted in reporter solution (Akoya Biosciences) to a 1:200 dilution in sets of three, as broken down in SI Table 1). The CODEX system automatically dispensed antibody solution and removed antibody solution automatically. Microscopy was performed on a Zeiss Axio Observer (Carl Zeiss AG, Oberkochen, Germany) using a Colibri 7 LED lightsource (Carl Zeiss) and C13440 camera (Hamamatsu, Shizuoka, Japan). Images were acquired with a 10% tile overlap and a z-stack ranging from 11 to 20 tiles (depending on the tissue size). Images were processed using the Akoya processor (Akoya biosciences) and QuPath.[1]

(1) Bankhead, P.; Loughrey, M. B.; Fernández, J. A.; Dombrowski, Y.; McArt, D. G.; Dunne, P. D.; McQuaid, S.; Gray, R. T.; Murray, L. J.; Coleman, H. G.; James, J. A.; Salto-Tellez, M.; Hamilton, P. W. *Scientific Reports* **2017**, *7*, 16878.
